# Protective immunity against severe malaria is associated with antibodies to a conserved repertoire of PfEMP1 variants

**DOI:** 10.1101/472159

**Authors:** Sofonias K. Tessema, Rie Nakajima, Algis Jasinskas, Stephanie L. Monk, Lea Lekieffre, Enmoore Lin, Benson Kiniboro, Carla Proietti, Peter Siba, Philip L. Felgner, Denise L. Doolan, Ivo Mueller, Alyssa E. Barry

## Abstract

Extreme diversity of the major surface antigen and virulence determinant of the malaria parasite *Plasmodium falciparum*, Erythrocyte Membrane Protein-1 (PfEMP1), poses a major barrier to identifying targets of protective immunity. To overcome this problem, we developed a PfEMP1 protein microarray containing 456 DBLα domains, which was used to characterize the immunome of a cohort of semi-immune children and to identify variants associated with protective immune responses. Children with high mean antibody levels to DBLα group 2 had a 26-36% reduced risk of uncomplicated (clinical) malaria, however only 8 diverse DBLα variants were weakly associated with protection from clinical malaria and had low predictive accuracy. On the other hand, children with high mean antibodies to DBLα groups 1 and 2 (which are markers for pathogenic “Type A” PfEMP1) and elevated antibodies to 85 (18.6%) of individual DBLα variants had a 70 −100% reduced risk of severe malaria. Of the top 20 predictive variants for severe disease protection, 17 were strongly associated with protection (86 - 100% reduction in risk of severe malaria) and had high predictive accuracy for severe disease risk. Many variants were conserved and had highly correlated antibody responses, including the three highest-ranking variants, which were linked to EPCR-binding CIDR domains. The results suggest that while immunity to uncomplicated malaria is characterised by antibodies to a diverse repertoire of PfEMP1, immunity to severe malaria requires antibodies to a limited subset of antigenically conserved variants. These findings provide new insights into antimalarial immunity and potential biomarkers for tracking disease risk.

## Introduction

Antigenic diversity of human pathogens is critical for evasion of host immune responses and is a significant challenge for the development of vaccines and other interventions. Human malaria parasites employ this strategy to achieve super-infection and to re-infect previously exposed hosts *(1, 2)*. In addition, they utilize clonal antigenic variation to persist within hosts and maximize transmission potential *(3)*. *Plasmodium falciparum* Erythrocyte Membrane Protein 1 (PfEMP1), expressed on the surface of infected erythrocytes, is the dominant target of naturally acquired immunity to malaria *(4)*. To evade immune responses, parasites switch PfEMP1 variants through differential expression of distinct members of the *var* multigene family *(5–7)*. PfEMP1 plays a key role in parasite virulence via cytoadhesion of infected erythrocytes to various host cell receptors *(8)*. Naturally acquired immunity develops with frequent exposure to *P. falciparum* infections, and is thought to require the piecemeal acquisition of antibodies to distinct PfEMP1 variants circulating in the parasite population *(9)*. This immunity is non-sterilizing and presents as a reduction in the severity of malaria symptoms with age *(9, 10)*. The acquisition of immunity to severe malaria early in life after only a small number of infections *(11)* suggests that clinically important PfEMP1 are antigenically conserved, whereas immunity against mild disease develops more slowly and may require a broader repertoire of antibodies to diverse PfEMP1 variants. Defining these immune targets is a key priority for the development of interventions to reduce the burden of malaria, including vaccines and biomarkers of exposure *(12)*.

A major hurdle to overcome in the identification of clinically important PfEMP1 variants is the extreme genetic diversity of the multicopy *var* genes that encode these antigens *(13–15)*, and a lack of appropriate tools to consider the full spectrum of diversity. Each *P. falciparum* genome harbors up to 60 distinct *var* genes with high levels of sequence diversity both within and between genomes *(16)*. The <u>D</u>uffy <u>B</u>inding <u>L</u>ike alpha (DBLα) domains present within the relatively conserved head structure of almost all *var* genes have been used as a marker for *var* gene diversity and gene expression studies *(14, 23–27)*. In order to extrapolate these findings to full length *var* genes with diverse functions, short (150-200 amino acid residue) DBLα sequence tags have been classified into six subgroups based on the number of cysteine residues (Cys2, Cys4, CysX) and specific sequence motifs (MFK and REY) in positions of limited variability (Cys/PoLV, *(17, 18))*, whilst full length DBLα domains have been classified into 3 subgroups (DBLα0, 1, 2) based on evolutionary relationships *(19)*. Group 1-3 DBLα tags (Cys2) are found in group A *var* genes and are expressed at high levels in parasites infecting young children with limited immunity and with severe disease *(20–22)*. DBLα groups 4-6 (Cys4/X) account for the majority of the sequences from non-group A *var* genes *(23)*. Expression of group 1 DBLα (Cys2 MFK+REY-) is associated with impaired consciousness and cerebral malaria *(21, 22)*, whilst expression of group 2 DBLα (Cys2 MFK-REY+) and group 5 DBLα (Cys4 MFK-REY+) has been associated with rosetting and severe malarial anaemia *(17, 24)*. Interestingly, groups 1 and 2 are also more conserved than other DBLα classes *(17)*. DBLα also plays a direct role in cytoadhesion *(25)* and anti-DBLα antibodies disrupt rosettes, reduce parasite cytoadhesion and are associated with clinical immunity in hyperendemic regions *(26, 27)*. Antibody responses to recombinant DBLα tags may therefore reveal exposure to associated full-length, clinically significant, *var* gene classes. Anti-DBLα antibodies also have potential as biomarkers of immunity against malaria and may directly confer protective immunity, however no link between antibodies to specific subgroups or individual variants and clinical or severe malaria has been made to date.

Clinical or severe malaria-associated PfEMP1 variants have potential as targets of malaria interventions, however identifying the most important variants amongst the thousands circulating in natural parasite populations *(13, 14, 28–30)* requires novel approaches. We sought to investigate antibody responses using parasite-encoded DBLα domains and human plasma derived from semi-immune children naturally exposed to these parasites, to address key questions about immunity to PfEMP1 including: i) which recombinant PfEMP1 domains are recognized broadly by naturally acquired antibodies, ii) are these antibodies protective against clinical and severe malaria, and iii) how conserved are key antibody targets *(12)*. In order to identify PfEMP1 variants that are key targets of immunity against severe and clinical disease, we developed a protein microarray containing 456 distinct DBLα variants to represent the PfEMP1 (*var* gene) diversity in the Papua New Guinea (PNG) parasite population *(14)*. Using this array, we seroprofiled PNG children aged 1-3 years old that were followed prospectively for clinical and severe malaria, allowing associations to be measured between the DBLα antibody repertoire and risk of disease. Candidate DBLα variants were then matched to identify corresponding full-length *var* genes composition in the PNG parasite population. The results provide key insights into the early acquisition of antibody responses to PfEMP1, identify biomarkers and potential targets of protection against clinical and severe malaria, and exposes links between the genetic and antigenic diversity of PfEMP1 that are critical to its development as a vaccine candidate.

## Results

### DBLα protein microarray design and quality control

We developed a protein microarray representing the *var* gene diversity of three parasite populations of PNG (Fig. S1). Validation of the protein array is described in the Materials and Methods and in the Supporting Material (Text S1, Fig. S2). After quality control, which included resequencing, successful cloning into the expression vector and checking for high quality protein spots on the array, antibody data from 456 DBLα sequences were included in the analyses (Text S1, Fig. S1, S2). This includes 42 group 1 (Cys2 MFK+REY-), 10 group 2 (Cys2 MFK-REY+), 32 group 3 (Cys2 MFK-REY-), 278 group 4 (Cys4 MFK-REY-), 63 group 5 (Cys4 MFK-REY+) and 31 group 6 (CysX, MFK-REY-) variants. This reflects the proportions found among 4656 full length DBLα domains extracted from 4656 *var* genes assembled using whole genome sequence (WGS) data from 122 independent PNG isolates as we described previously *((31)*, Fig. S1). Group 2 DBLα variants were underrepresented on the protein microarray relative to the WGS data (2.2% vs 4.7%, *p* = 0.008) (Fig. 1A), which may be due to primer bias or the lower diversity of this subgroup *(17)*.

**Fig. 1.**
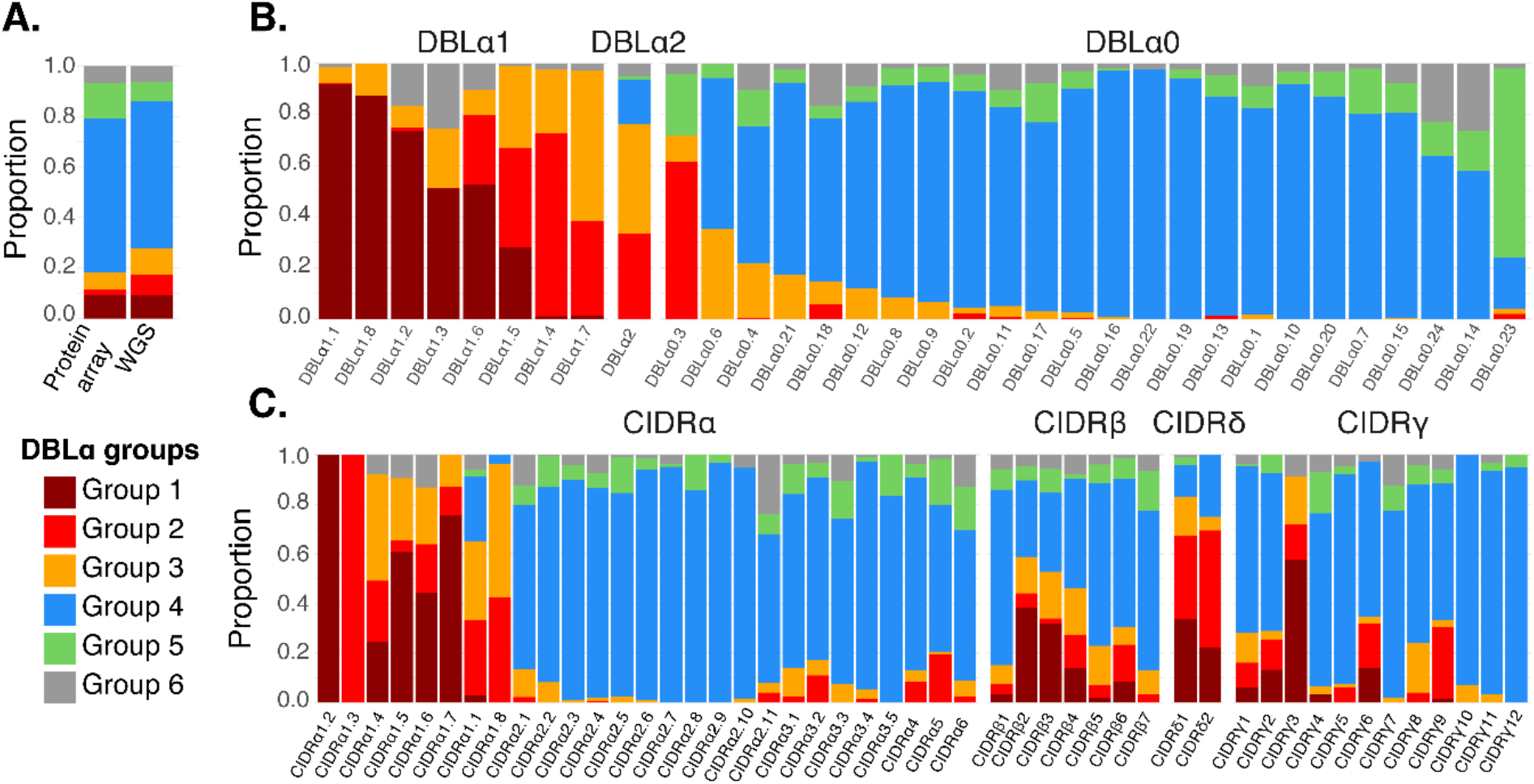
Relationship between DBLα tags and full length DBLα and CIDR domain classes in PNG. (**A**) The proportion of the six DBLα tag classifications were compared for the 456 DBLα variants and for 4656 DBLα domains extracted from full length *var* genes sequenced from 122 WGS of Papua New Guinean isolates. (**B**) Relationship between DBLα tags and full length domains classified according to Bull *et al*. 2007 (*17*) and Rask *et al.*, 2010 (*19*) respectively for 4656 DBLα domains extracted from 122 PNG isolates. (**C**) Relationship between DBLα tags and full length CIDR domain classifications in 6142 CIDR domains extracted from the same *var* genes in 122 PNG isolates.

To further evaluate the diversity of DBLα using the PNG WGS data, we compared the distribution of DBLα classes among the alternative DBLα classes *(19)* and association with CIDR domain classes. Of note, groups 1, 2 and 3 DBLα variants were found within DBLα1.1-1.8 domains, which are associated with the upsA promoter of type A *var* genes *(19, 32)*. These groups are also associated with Endothelial Protein C Receptor (EPCR) binding (e.g. CIDR1) and rosetting-associated CIDR domains (CIDRb/d/g) *(32)*. In addition, DBLα2, a type B *var* gene domain associated with CIDRα1.1/1.8 in DC8 head structures *(19)*, were largely composed of group 2 (33%) and 3 (43%) DBLα variants (Fig. 1). Therefore, the developed protein microarray is the most comprehensive protein array to date representing the genetic and functional diversity of DBLα domains found in naturally circulating parasite populations.

### Repeated exposure drives the acquisition of antibody responses to DBLα in young children

Baseline plasma samples from a longitudinal cohort of 232 young children (median age 1.7 years, range 0.9 to 3.2 years) from the Ilaita region, East Sepik Province, PNG ((*33*), Fig. S1), were screened for antibody responses (total IgG) to the 456 DBLα variants on the microarray. Pooled plasma from naïve Australian adults (n=20) showed limited reactivity, PNG children (aged 1-3, n=25) showed moderate reactivity, and PNG adults (n=20) had overall high reactivity to the different DBLα variants (Fig. 2A, Fig S3). Antibody profiles of individual children showed differential recognition of DBLα variants (Fig. 2B). By stratifying the antibody response data according to the six DBLα subgroups, we found that group 1 and group 2 variants were more frequently recognized (*p* < 0.0001), while antibody levels to groups 3-6 were generally low (Fig. 2C). The sero-dominance of group 1 and 2 variants was evident across all ages in the cohort (Fig. 2D).

**Fig. 2.**
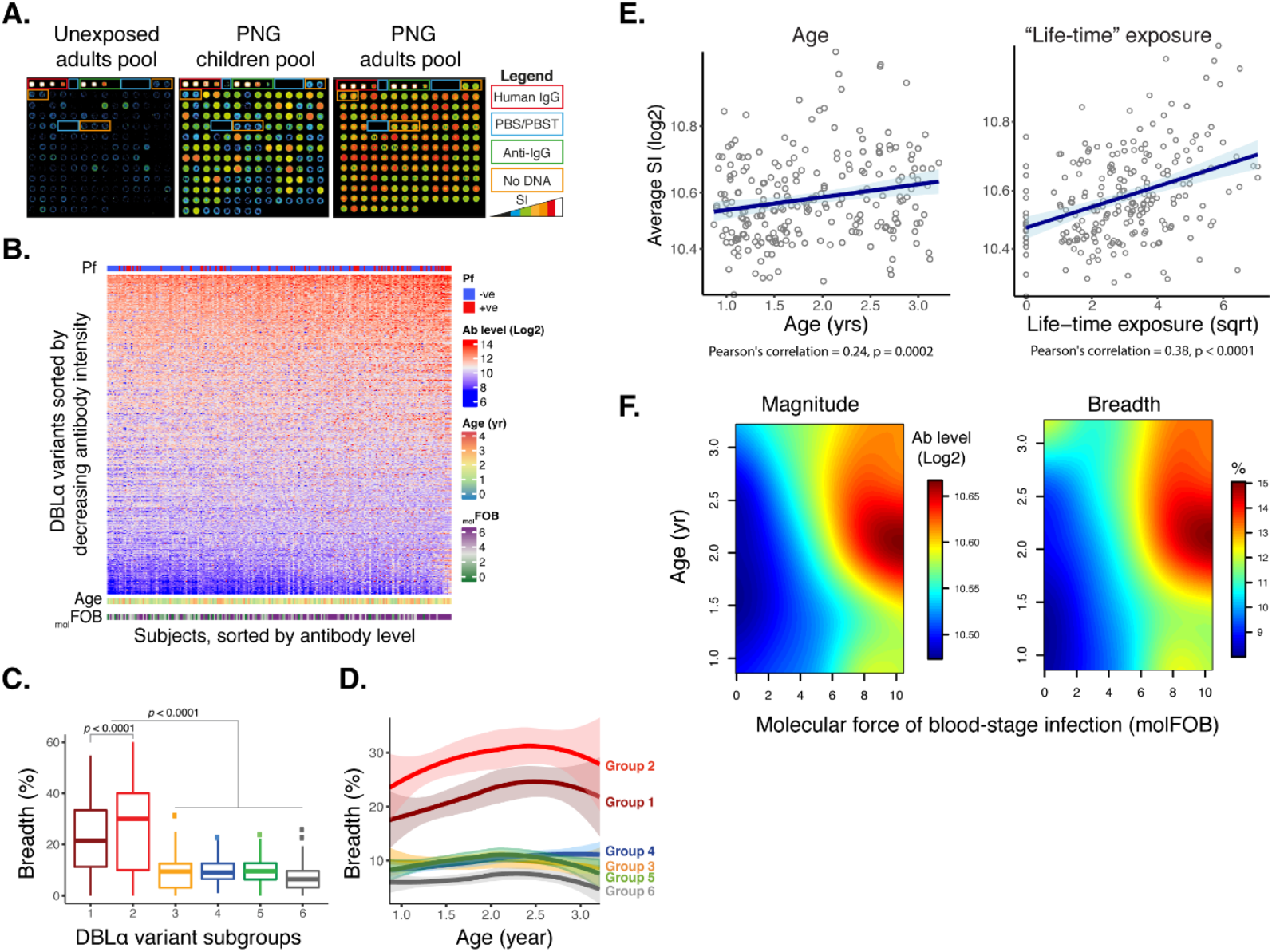
Antibody responses to 456 DBLα variants for 232 young children. (**A**) Representative images of control experiments. Protein arrays probed with pooled negative and positive control (children and adults) plasma samples. Each spot represents a protein antigen or control protein (coloured boxes). Reactivity is presented as Signal Intensity (SI), with spots saturated at ~65,000 (white colour). The overall responses are quantified and summarized for 456 variants in Fig. S3. (**B**) Heat map showing signal intensity (log2) to all 456 DBLα variants show that responses are highly variant specific. Children were ordered by the average magnitude of antibody intensity. The infection status, age and molecular force of blood stage infection (_mol_FOB) of the children are indicated at the top and bottom bars, respectively. (**C**) Breadth of antibody response to six DBLα subgroups. The proportion of antigens recognised for each of the 232 children is presented. (**D**) Smoothed (local weighted regression) breadth of antibody response to six DBLα subgroups with age. (**E**) Correlation of magnitude of average antibody levels with age and life-time exposure. (**F**) Magnitude and breadth of antibody responses to DBLα variants as a function of age and individual exposure as measured by _mol_FOB.

We hypothesized that if exposure to the DBLα variants circulating in an endemic area increases with age and repeated exposure to infection, then the overall antibody response will increase in magnitude (mean antibody titre) and breadth (mean number of antigens recognized). In an initial exploratory analysis, DBLα antibodies were associated with age (antibody level, *p* = 0.014; breadth, *p* = 0.002) and concurrent infection (*p* < 0.0001), as well as average exposure as predicted by the number of infections during the follow-up period (molecular force of blood stage infection, _mol_FOB, *p* < 0.0001, Fig S5) *(34)*. Overall antibody level significantly increased with age (Pearson’s r = 0.24, *p* = 0.0002), _mol_FOB (Pearson’s r = 0.29, *p* < 0.0001) and “life-time” exposure (age scaled by _mol_FOB, Pearson’s r = 0.38, *p* < 0.0001) (Fig. 2E).

Due to the complex interaction of age and _mol_FOB with magnitude and breadth of antibody response (Fig. 2F), we assessed the explanatory power of each predictor in a linear regression model. In a univariate analysis, both age (estimate = 1.38, *p* = 0.0003, R^2^= 0.051) and _mol_FOB (estimate = 0.57, *p* < 0.0001, R^2^=0.13) are significantly associated with breadth of responses with _mol_FOB explaining the variability better than age. In a multivariate analysis, both age and _mol_FOB were significantly associated with breadth (Age_estimate_ = 0.88, *p* = 0.02; _mol_FOB estimate, = 0.51, *p* < 0.0001, R^2^=0.15). We then determined the relative contribution of the predictors, using an averaging of the sequential sum-of-squares, considering both direct effect (i.e. correlation) and effects when combined in the regression equation. With this approach, _mol_FOB explained 75% [95% CI: 48 – 93.5] of the variability in the breadth of antibody response, whereas age only 25% [95% CI: 6.5 – 52], which suggests that in these children _mol_FOB explains the differences in the antibody levels more accurately than age. Thus, we are in a unique position to accurately assess the association of antibody responses with prospective risk of severe and clinical malaria after correcting each child’s antibody response for the key confounders of age and _mol_FOB.

### Antibodies to a subset of DBLα variants are associated with reduced risk of clinical malaria

Given that different PfEMP1 subgroups are expressed during clinical, uncomplicated malaria (*22, 23*), high levels of antibodies against virulence-associated variants may also be associated with reduced risk of uncomplicated malaria. Only 16 of the 456 variants were associated with a 30-40% reduction in the risk of clinical malaria, and 14 variants were associated with a 33-45% reduction in the risk of high-density clinical malaria. Eight variants were associated with reduced risk of both low and high-density clinical malaria (Fig. 3A). Group 5 DBLα variants were overrepresented among those associated with protection from clinical (7/16, *p* < 0.003, exact binomial test) and high-density clinical malaria (6/14, *p* < 0.008). When antibody levels were averaged for DBLα sub-groups, we found that children that were high responders to group 2 DBLα variants had a 26% reduction in the incidence of all categories of uncomplicated clinical malaria (aIRR = 0.74, 95% CI: 0.56-0.97; *p =* 0.028) and a 36% reduction in high-density clinical malaria (aIRR = 0.64, 95% CI: 0.47-0.89; *p* = 0.008) as compared with low-responders (Fig. 3B). In contrast, high responders to group 4 DBLα variants had a 43% increased risk of clinical malaria, suggesting that group 4 variants are indicators of risk of exposure rather than disease protection.

**Fig. 3.**
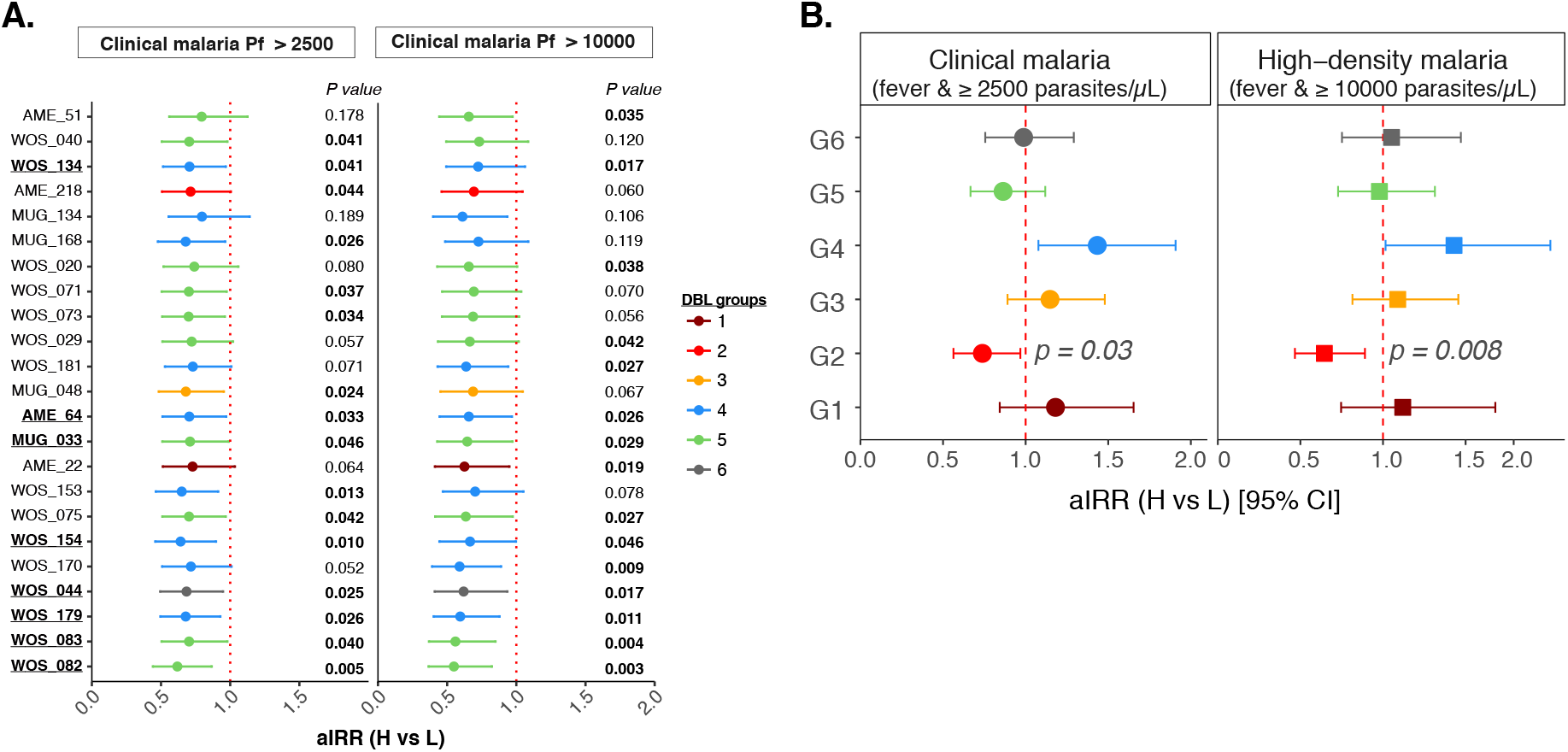
Association of antibodies to six DBLα groups with protection against clinical malaria. (**A**) The incidence rate of clinical malaria and high-density clinical malaria was compared for high and low tertiles based on antibody response (MFI) to each group of DBLα variants. The regression model was adjusted for age at enrolment, *P. falciparum* infection status and differences in individual exposure (_mol_FOI) and village of residence. The adjusted incidence rate ratio (aIRR) for the comparison of high and low responders and the 95% confidence intervals for only significant variants are shown. Variants that are significantly associated with protection from clinical and high-density clinical malaria are bold and underlined. ROC curve generated for the top 20 variables (**B**) and the six groups of DBLα variants

### Antibodies to specific DBLα variants are associated with reduced risk of severe malaria

We then investigated the association of anti-DBLα antibodies with the prospective risk of severe malaria in the cohort. Of the 232 children, 19 (8.2%) had severe *P. falciparum* malaria during the follow-up period *(33)* according to WHO criteria *(35)*, though they were similar to the remaining 213 children with respect to age (1.58 years, range 0.98 to 2.96 versus 1.88 years, range 0.87 to 3.2 years, respectively) and exposure (_mol_FOI = 5.34 ± 3.4 versus 5.71 ± 4.4 respectively). Associations between antibody levels at enrolment and the occurrence of severe malaria during the follow-up period were then assessed for each DBLα variant by determining the incidence rate ratio of severe malaria for high versus low responders, adjusted for confounders (aIRR). Eighty five (85/456, 18.6%) DBLα variants were significantly associated with protection (70 to 100% reduced risk) of severe malaria in high responders compared to low responders (Fig. 4A, Table S1). The DBLα subgroups associated predominantly with type A *var* genes were significantly over represented among the protective variants (group 1: 9% on the array vs 30% of the protective variants, *p* < 0.001, binomial test; and group 2: 2% vs 5.9%, *p* = 0.03, binomial test). In contrast, group 4 variants represent 61% of the variants on the array and only 26% of the protective variants, suggesting antibodies to this subgroup play a minor role in protection against severe malaria.

**Fig. 4.**
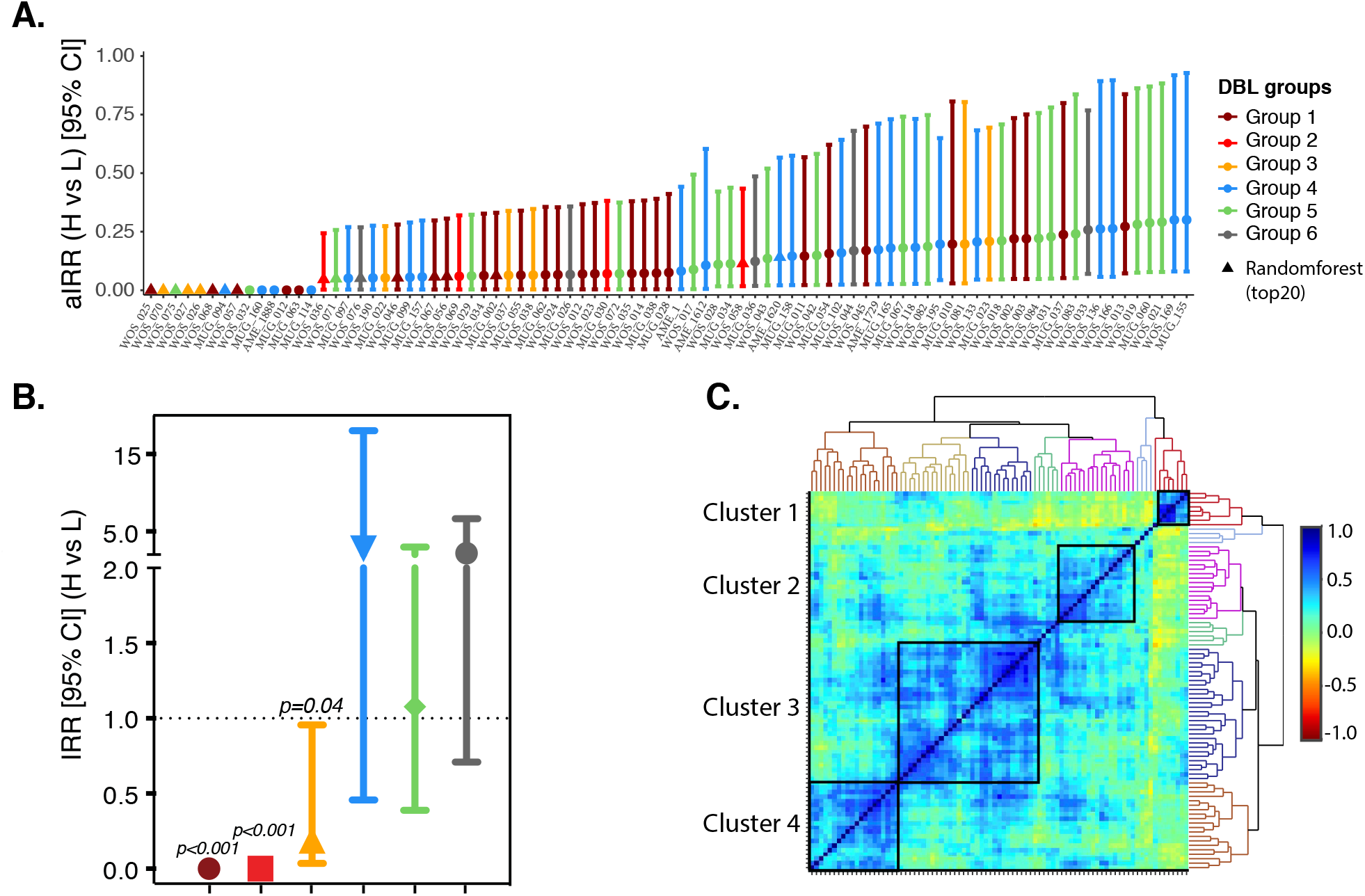
Antibody levels to DBLα variants and severe malaria. (**A**) Association between antibody levels to DBLα variants and prospective risk of severe malaria. Individual responses were grouped into tertiles. The incidence rates of severe malaria were compared for high and low responders for each DBLα variant. The regression model was adjusted for age at enrolment, *P. falciparum* infection status and differences in individual exposure (_mol_FOI) and village of residence. The adjusted incidence rate ratio (aIRR) for the comparison of high and low responders and the 95% confidence intervals for only significant variants are shown. The p-values for the associations are attached in supplementary Table S1. (**B**) Association of antibodies to six DBLα groups with protection against severe malaria. Individuals were grouped into tertiles based on average MFI to each DBLα sub-group. The incidence rate of severe malaria was compared for high and low responders for each subgroup of DBLα variants. Similar adjustments were made in the regression model as panel A. The p-values are indicated only for significant (p < 0.05) values. (**C**) Heat map showing the pairwise correlation coefficients of antibody responses to protective DBLα variants stratified by clusters (solid lines).

Antibody levels averaged for the six DBLα subgroups showed high responders to group 1 and group 2 DBLα variants had a 100% reduced risk compared to low responders (Fig. 4B), suggesting a combined repertoire of antibodies to these groups in early childhood are protective against severe malaria. Consistent with this finding, the majority (73%) of the 3570 pairwise correlations were significantly positively correlated (rho = 0.3 [range: 0.13 - 0.83], *p* < 0.05) (Fig. 4C). Clustering analyses of the correlation coefficient identified four positively correlated clusters, cluster 1 is exclusively composed of group 1 DBLα variants (n=7), whereas all the other clusters are composed of multiple groups, indicating high levels of antibody co-acquisition and/or cross-reactivity among protective variants (Fig. 4C). A neighbour joining tree shows that the majority of the protective variants are from a relatively conserved clade, which correspond to variants from type “A” *var* genes *(13)*, however there were also diverse protective variants outside this clade (Fig. S5).

### Children with severe malaria during the follow-up period acquire antibodies to protective variants

We then explored the dynamics of antibody responses over time by comparing antibody levels at enrolment and at the end of the 16 months of the follow-up period. At enrolment, severe disease cases had lower levels of antibodies overall and to protective variants, compared to the non-severe cases (*p* < 0.0001), but there was no significant difference in antibody responses to non-protective variants. By the end of the follow-up period, after an average of 6 six infections per child, antibody levels to all variants and specifically to those protective variants were boosted (*p* < 0.0001). No significant difference in antibody response was observed in non-severe cases for the protective variants, but there was a significant increase for the non-protective variants (*p* = 0.0002) (Fig. 5).

**Fig. 5.**
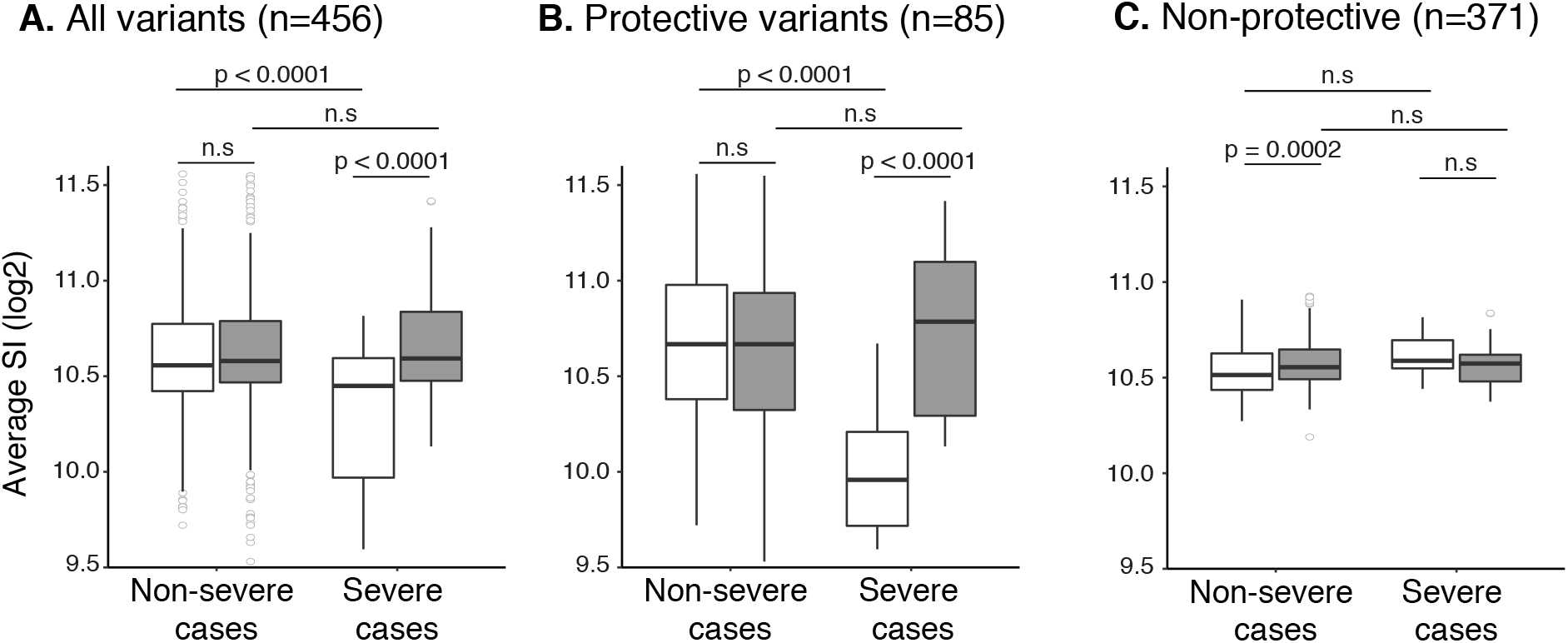
Dynamics of antibody responses with severe disease. Antibody levels are shown for severe and non-severe malaria cases at enrolment (open boxes) and at the end of 16 months of follow-up period (grey boxes) for (**A**) all variants (**B**) protective and (**C**) non-protective variants. Bonferroni adjusted p-values are shown at the top of each pairwise comparison.

### Antibodies to protective DBLα variants predict prospective risk of severe but not clinical malaria

The lack of antibody responses to protective variants in young children who developed severe malaria and boosting of these antibody responses by the end of the follow-up period suggests that certain PfEMP1 antibodies may be a useful predictor for future risk of severe malaria. To test this hypothesis, we built a data driven machine-learning model to identify predictive variants. The top 20 variants, based on the mean decrease in classifier accuracy, were selected for further analyses (Table S1). Of these, 17 were significantly associated with protection (86 - 100% reduction in risk of severe malaria) in the analysis above, therefore we focused all further analysis on these variants (Table S2). The 17 variants included representatives from all 6 DBLα sub-groups, however group 1 variants were dominant (41%, p<0.0001, exact binomial test). Furthermore, at enrolment, antibody levels to the 17 variants were significantly lower in severe cases compared to non-severe cases (*p* < 0.0001, Fig. S6), and were boosted by the end of the follow-up period (Fig.S6), despite the lack of difference in force of infection between the two groups.

In order to evaluate the predictive utility of antibodies in discriminating between children who go on to experience severe malaria and those who do not, receiver operating characteristic curves (ROC) were generated and the area under the curves (AUC) determined. The AUC is associated with high predictive accuracy when its value is close to 1. Antibodies against the top 17 protective and predictive DBLα variants had high predictive accuracy (median AUC = 0.82 [range: 0.66 - 0.88]) (Fig. 6A, Table S2). Three variants, WOS_026 (sub-groups 3 and DBLα2), WOS_025 (sub-groups 1 and DBLα1.1) and WOS_068 (sub-groups 1 and DBLα1.1) variants, had the highest AUC 0.88, 0.86 and 0.83 and are associated with 100% reduction in risk of severe malaria (Table S2). AME_1620 (sub-groups 4 and DBLα0.2) variant, had the lowest AUC (0.66) and the lowest incidence rate ratio (aIRR = 0.17) of the top 17 DBLα variants (Table S2). When averaged by DBLα subgroup, group 1 and group 2 DBLα variants had high predictive values (AUC ≥ 0.8), while all the other groups had low discriminatory power for prediction of severe malaria (Fig. 6B). This demonstrates that low antibody levels against DBLα groups 1 and 2 are predictive of prospective risk of severe malaria. However, antibody levels to any of the individual DBLα variants or subgroups had a poor predictive power for clinical malaria (AUC= 0.5 - 0.57). In fact, _mol_FOB was the major predictor of clinical malaria (AUC=0.8) but this was not the case for severe malaria (AUC= 0.49), and is consistent with previous findings (*34*) (Fig. S7).

**Fig. 6.**
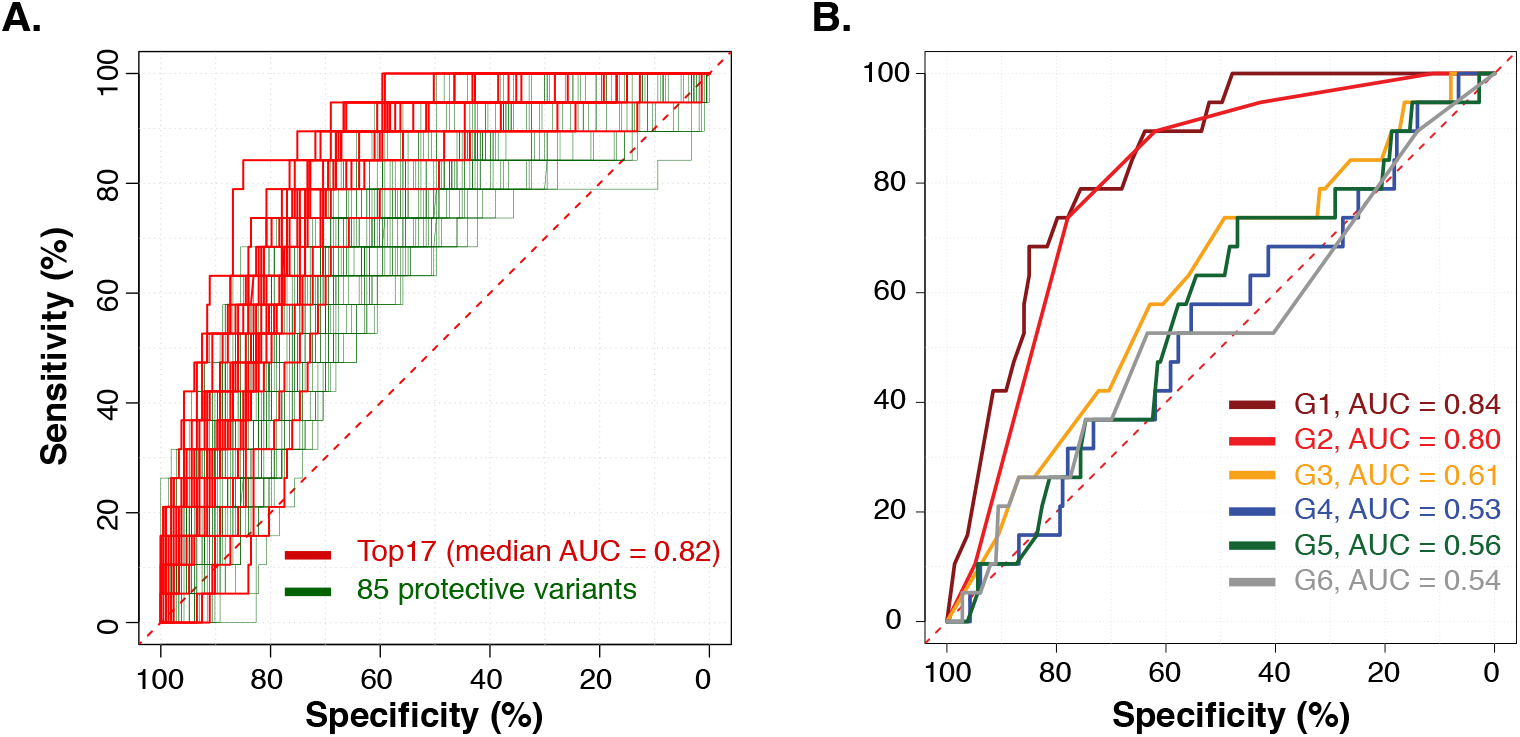
Antibody levels to DBLα variants and prediction of risk of severe malaria. Receiver-Operating Characteristic (ROC) curve generated for (**A**) the 85 protective and (**B**) the six groups of DBLα variants. The Area Under the ROC Curves (AUC) are shown for the top 17 in panel A and the six DBLα groups in panel B. AUC values for the top 17 highly predictive variants are given in Table S2. The red dashed diagonal line represents the ROC curve for a test with no discriminatory ability (AUC=0.5).

## Discussion

The diversity of PfEMP1 poses a formidable obstacle to the study of naturally acquired immunity to malaria and to the rational design of vaccines targeting this antigen. Our results show that semi-immune PNG children broadly recognize recombinant DBLα domain variants with stronger antibody response to group 1 and 2 variants. Protection against clinical malaria was associated with high antibodies to only 16 individual DBLα variants, with a significantly enriched proportion of group 5 variants. When averaged by DBLα group, group 2 variants were associated with reduced risk of clinical malaria. However the prospective risk of clinical malaria was more accurately predicted by molecular force of blood stage infection (_mol_FOB) than variant-specific antibody responses. Protection against severe malaria however was strongly associated (up to 100% reduced risk) with high mean antibodies to both group 1 and 2 DBLα and 85 individual variants (enriched for group 1 and 2 DBLα variants). At the end of the follow-up period antibodies to these 85 severe-disease protective variants were significantly higher, indicating that these antibodies were acquired through natural infection as the children were exposed in average to six new infection during the follow-up period. Furthermore, we identified the top 20 highly predictive variants through machine learning, revealing that low antibodies to 17 of the 85 protective variants were also strong predictors of the prospective risk of severe malaria. Many of these antigens showed highly correlated antibody responses, and the 3 highest-ranking variants were associated with EPCR binding CIDR domains. These results suggest that whilst a large and diverse repertoire of antibody responses are required for protection against clinical malaria, antibody responses against a handful of conserved variants may be sufficient for protection against severe malaria.

Children develop protective antibodies specific to the infecting parasite isolate following infection *(4)* and these antibodies are thought to provide subsequent protection from heterologous isolates *(4, 36–39)*. Associations of antibodies to specific PfEMP1 variants with protection from severe and clinical malaria could be obscured by the confounding effects of cumulative exposure due to the high diversity and immunogenicity of PfEMP1 *(12)*. In this study, we were able to adjust for the confounding effects of individual exposure (i.e. _mol_FOB), age, infection status and village of residence. Eighty five (70-100% reduced risk) and 16 (30-40% reduced risk) of the 456 DBLα variants were significantly associated with protection from severe and clinical malaria, respectively. The larger proportion and stronger association of variants associated with severe malaria supports the notion that severe malaria is caused by the expression of a relatively conserved subset of PfEMP1 whilst protection against clinical malaria may require antibodies to a larger diversity of PfEMP1. When aggregated by DBLα group, none of the children with high levels of antibodies against group 1 and 2 variants developed severe malaria (i.e. 100% reduced risk). In addition, antibodies against group 3 DBLα variants were associated with an 83% reduced risk of severe malaria. This is the first study that has shown a strong association of antibody responses with specific PfEMP1 variants and protection against severe malaria. This implies that PfEMP1 containing these DBLα subgroups are important targets (or co-expressed with important targets), of immunity against severe malaria.

Exploring the dynamics of variant-specific antibody responses in the children who experienced severe malaria revealed significantly lower levels of antibodies to all DBLα groups compared to other children, suggesting that low levels of PfEMP1 antibodies in general may be associated with susceptibility. The strong associations of antibodies to DBLα groups 1, 2 and 3 (Cys2) and protection from severe malaria suggest that children with high levels of antibodies to these groups may be able to control disease and are protected from severe malaria. This is supported by the negative correlation of expression of these groups with anti-VSA antibodies in another study *(18)*. Children with severe malaria had significantly lower levels of antibodies to the protective variants before the onset of severe malaria. These antibodies were significantly boosted during the severe malaria episode and remained high in the convalescent period. Our analysis of the predictive accuracy of low antibodies to group 1 and 2 DBLα variants, as well as the top 17 predictive and protective variants for severe malaria, suggests they may be developed as biomarkers for antimalarial immunity against severe disease. These biomarkers would be extremely useful to identify high-risk human populations in regions attempting to eliminate malaria.

Our results suggest that a broad repertoire of antibodies is important for protection against clinical malaria. Nevertheless, mean antibodies to group 2 DBLα variants reduced the risk of clinical, and high-density clinical malaria. Previous studies showed that elevated expression of group 2 variants is associated with rosetting and severe malaria *(18, 20–22, 28, 40)*. In addition, we previously found group 2 to be serodominant in PNG infants less than one year of age, consistent with these variants being expressed in early infections *(13)*. The association of group 2 antibodies with a reduced risk of clinical episodes represents a major advance in defining and limiting the subset of PfEMP1 containing these domains as functional immune targets and warrants further investigation, because group 2 variants account only for 2 to 10% of the total DBLα variants sampled so far in several geographic areas *(13, 14, 17, 20, 29, 40-42)*.

There is a strong rationale for a PfEMP1 vaccine that would protect against clinical and severe malaria, since any reduction in disease and death would ease the huge burden of malaria infection on susceptible individuals *(43)*. Leading candidates to protect against severe malaria include specific classes of PfEMP1 variants and domain cassettes (DCs) that have been linked to the pathogenesis of severe malaria through *in vivo* and *in vitro* expression studies *(12)*. These include VAR2CSA [also known as DC2 (*44*)], DC4, DC8 and DC13 *(45–49)*, which are shared by majority of the *P. falciparum* clones analyzed *(20)* and exhibit conserved features linked to cytoadhesion *(50)*. Recent studies have also demonstrated functional and structural conservation of the highly diverse CIDR domain *(51-53)*, however no conserved epitopes have yet been identified in naturally circulating parasite populations. In the same cohort studied here, antibodies to ICAM-1 binding DBLβ domains were associated with protection against clinical malaria and the predicted ICAM1-binding DBLβ PfEMP1 were enriched for DC4 and DC13 head-structures, previously implicated in severe malaria *(31)*. In the current study, analysis of full-length *var* gene sequences from PNG isolates showed group 1 and 2 DBLα variants were strongly associated with DC4 and DC13 type CIDR domains. As these variants were also associated with protection, this suggests conservation of these domain cassettes and multiple binding specificities *(52. 53)*. Despite the extreme global diversity of *var* genes, the data presented herein links antibody responses against PfEMP1 variants previously implicated in severe malaria with strong protection from severe malaria. These variants are overrepresented by group 1 variants, the most conserved DBLα variants found exclusively in Type A *var* genes (17). To pursue PfEMP1 antigens as viable vaccine candidates, it will be critical to define the antigenic diversity of leading candidates, and to assess whether immune responses to geographically diverse PfEMP1s are associated with protection from clinical and severe malaria in multiple endemic areas. The use of high-throughput technologies, such as protein microarrays that we have employed here, together with well-designed cohort studies will be essential for this task.

We have demonstrated that antibodies to limited repertoires of DBLα variants representing a fraction of the overall PfEMP1 diversity are acquired early in life, and are significantly associated with a reduced risk of clinical and severe malaria. Antibodies against a subset of these variants predict a lower risk of severe malaria with high specificity and sensitivity providing support for their use as biomarkers of protection and supporting their development as vaccine candidates. The development of any vaccine based on pathogenic PfEMP1 will require further functional characterization, as well as analysis to identify conserved variants that are able to induce variant transcending protective immune responses across different geographic regions.

## Materials and Methods

### Ethics Statement

This study was approved by the Institutional Review Board of the PNG Institute of Medical Research (10.21), the Medical Research and Advisory Committee (MRAC) of the Ministry of Health in PNG (10.55) and the Human Research Ethics Committee of the Walter and Eliza Hall Institute (11.03).

### Study Area and Samples

The PNG north coast is an area of perennial high transmission of *P. falciparum* malaria *(53)*. Plasma samples were collected in a longitudinal cohort survey conducted in Ilaita area in the East Sepik Province of PNG in 2006 *(33)*. Briefly, 264 children aged 1–3 years of age (median 1.70 years) were enrolled and followed for 69 weeks. The study consisted of three fortnightly surveillance visits and each concluding with the collection of two blood samples 24 hours apart for active detection of malaria infection. Incidence of clinical malaria in each 8–9 week follow-up interval was estimated as previously described *(31, 33)*. Clinical malaria was defined as febrile illness (axillary temperature ≥37.5°C or history of fever in preceding 48 hrs) with a concurrent *P. falciparum* parasitemia >2500 parasites/μl. High-density clinical malaria was defined as febrile illness with a concurrent *P. falciparum* parasitemia >10000 parasites/μl. Infection and _mol_FOB was determined using light microscopy and PCR *(31, 33)*. The DBLα variants were amplified from genomic DNA collected from three geographical areas of PNG, namely Wosera (East Sepik Province) in 2005, Mugil (Madang Province) in 2006 and Amele (Madang Province) in 1999 (Fig. S1). The details of these studies are published elsewhere *(13, 14)*.

### Protein microarray production, validation and serological screening

Protein microarray chips were constructed as described previously *(56–61)*. Protein arrays were used to probe 3μl of plasma samples diluted in 1:100 protein array-blocking buffer (Maine Manufacturing) with 20% (w/v) *E. coli* lysate, which significantly reduced the background as compared to the conventional 10% (w/v) *E. coli* lysate (*p* < 0.0001, Wilcoxon rank sum test) (Fig. S2A). Therefore, in all assays 20% (w/v) *E. coli* lysate in blocking buffer was used. Antibody assays were performed using paired plasma samples collected at enrolment and end of the follow-up from 232 children. The assays were done as described previously *(56–61)*. The proportion of DBLα groups represented on the protein array and the relationship between DBL and CIDR domains groupings were compared in 4656 DBLα and 6142 CIDR sequences obtained from 122 whole genome PNG sequences *(31)*.

### Data processing and normalization

Signal intensities were corrected for spot-specific background, where the per-spot local background was subtracted. The SI (Signal Intensity) was normalized against the “No DNA” controls using the variance stabilization and normalization method and using the VSN package installed in R-project *(62)*. The vsn-normalized data was used for all statistical analyses and visualization (mean intensity in log2).

The antibody responses to the DBLα variants were quantified using three measurements: magnitude, breadth and seroprevalence. Magnitude is the mean fluorescence intensity (MFI) for each DBLα variants on the array or the average intensity for a group of DBLα variants. Seroprevalence is determined by calculating the percentage of donors seropositive for each DBLα variant. Seropositivity to DBLα variants was determined by having signal intensity greater than 2 standard deviations above the average intensity of 20 malaria naïve Australian adult controls included in the screening. Breadth of response was determined by calculating the number of seropositive DBLα variants per child as a function of the total number tested (% DBLα variants recognized). The DBLα variants on the array were grouped in to six (Group 1 to 6) as described previously *(17)*. The average magnitude, breadth and seroprevalence were calculated according to these classifications.

### Statistical analysis

Statistical analysis of data was performed using Stata version 12.1 statistical software (Stata Corporation, Texas, USA) and R 3.2.3. Malariometric data from the cohort and antibody data were merged. Differences in the magnitude of antibodies with age (linear) and force of infection were assessed by using Pearson′s correlation tests. Differences in the median fluorescence intensity (MFI) between groups were compared using Kruskal-Wallis tests or Mann-Whitney U tests where appropriate. Association between age, molecular force of blood stage infection and magnitude or breadth of antibody responses was assessed using linear regression and relative importance was determined using the R package ‘relaimpo’ using the R^2^ partitioned by averaging over orders and the results were bootstrapped to determine 95% confidence intervals as described previously *(63)*. The exact binomial test was used to test for a biased distribution of subgroups amongst protective antigens.

A negative binomial GEE model (based on XTNBREG procedure in STATA 12.0) was used for the analysis of incidence of clinical and severe *P. falciparum* malaria. Two different models were used to assess the association of antibodies (for each individual variants and mean of the six groups) with the prospective risk of clinical malaria: i) ‘Unadjusted’ (IRR): no adjustment for confounding factors and ii) Adjusted for confounders (adjusted IRR): adjustment was made for all predictors of incidence of clinical malaria in this study. The model was adjusted for infection status at the time of antibody measurement (by light microscopy), seasonal variation (month, year) and spatial variation (village of residence), as well as for individual differences in exposure (_mol_FOI) and age (as a correlate of overall immune status). Only results from the adjusted models are presented. For DBLα variants that were significantly associated with protection, spearman’s rank test was conducted in order to determine correlation of antibody responses.

The R package ‘randomForest’ was used to build random forest binary classifiers in order to assess antibodies to which variants predict the prospective risk of severe or clinical malaria *(64)*. We used out-of-bag (OOB) error rate to measure the performance of the model. Out-of-bag error rate stabilized at 2,000 trees, so we chose this as a parameter for optimizing the number of variants sampled per node, which was determined as mtry = 3. The predictor variants were then evaluated based on the variable of importance. These models included antibody levels to each DBLα variant, age, infection status at the time of antibody measurement, molecular force of infections. These analyses were separately conducted for the two outcomes (clinical and severe malaria). Based on the mean decrease in accuracy (i.e. variable of importance measure), the top 20 variants were selected. For severe malaria, 17 of the top 20 variants were also significantly associated with protection from the previous analyses, therefore these 17 variants were evaluated for their prediction accuracy. For clinical malaria, molecular force of infection, age and infection status were ranked in the top 3 based on the variable of importance measures. Receiving Operating Characteristic (ROC) curves were used to estimate the prediction values of antibody levels at baseline whether an individual is susceptible for severe or clinical malaria during the follow-up period. For each ROC curve, the area under the curve and Youden index were calculated using the pROC package using R project *(65)*. These analyses were conducted for the 85 protective variants, top 17 predictors from the random forest and antibody responses aggregated by 6 groups. For clinical malaria, similar analyses were conducted for the top 20 variants from the random forest plus molecular force of infection and antibody responses averaged for the six DBLα groups.

## Supporting Materials

### Materials and Methods

**Text S1.** Validation of the protein microarray

### Results

**Table S1.** List of 85 protective variants with incidence rate ratio and p-values. *Top 20 variants from random forest binary classifier analyses are shown. 3 variants were not associated with protection from clinical malaria. Adjusted IRR as described in figure 4 are shown.

**Table S2.** Top 17 protective and predictive DBLα variants

**Fig. S1.** Details of the study sites, plasma samples and DBLα variants

**Fig. S2.** Validation and reproducibility of the protein microarray

**Fig. S3.** Recognition of DBLα variants by pooled control samples

**Fig. S4.** Magnitude and breadth of antibody responses to all variants by age, infection status at enrolment and exposure (_mol_FOB)

**Fig. S5.** Phylogenetic relationships of DBLα variants

**Fig. S6.** Acquisition of DBLα antibodies during the follow-up period

**Fig. S7.** Antibody levels to DBLα variants and prediction of clinical malaria risk

## Acknowledgements

We thank all study participants and their parents and guardians and all field staff at the Papua New Guinea Institute of Medical Research. In addition, we are grateful to T. Lavstsen, B. Petersen and J. Jespersen for *var* gene assembly as outlined in (*31*).

## Author contributions

AEB, IM and DLD conceived the study, supervised the experiments and contributed to the writing of the manuscript. SKT generated reagents, conducted the majority of experiments, analysed the data and wrote the manuscript, SLM generated reagents, PLF, RN and AJ produced the protein arrays and contributed to the interpretation of data, LL, CP and DLD supervised the antibody screen and assisted with data analysis, EL, BK and PMS collected samples and provided logistical support for the field studies.

## Data and materials availability

Antibody data and associated metadata is available upon request.

